# Dystroglycan proteolysis is conformationally-regulated and disrupted by disease-associated mutations

**DOI:** 10.1101/279315

**Authors:** Amanda N. Hayward, Wendy R. Gordon

## Abstract

The adhesion receptor dystroglycan provides a critical mechanical link between the extracellular matrix (ECM) and the actin cytoskeleton to help muscle cells withstand contraction and neural cells maintain the blood brain barrier. Disrupting the link is associated with diseases such as cancer and muscular dystrophy. Proteolysis of dystroglycan by Matrix Metalloproteases (MMPs) also breaks the mechanical anchor and is amplified in several pathogenic states. We use a combination of biochemical and cell-based assays to show that dystroglycan proteolysis is conformationally regulated by an extracellular, juxtamembrane “proteolysis domain”, comprised of tandem Ig-like and SEA-like domains. The intact proteolysis domain is resistant to MMP cleavage, but structurally-disruptive muscular dystrophy-related mutations sensitize dystroglycan to proteolysis. Moreover, increased dystroglycan proteolysis correlates with faster cell migration, linking proteolysis to a disease-relevant cellular phenotype. Intriguingly, previously uncharacterized cancer-associated mutations that map to the proteolysis domain similarly lead to increases in proteolysis and rates of cell migration, potentially revealing a new pathogenic mechanism in cancer.

## Introduction

Dystroglycan is an extracellular matrix (ECM) receptor that is typically expressed on the surface of cells that adjoin basement membranes, such as epithelial, neural, and muscle cells and plays important roles during development and adult homeostasis (1). Dystroglycan’s most well-understood function in cells is to provide a mechanical link between the extracellular matrix and the cytoskeleton (2). In its function as a core member of the dystrophin-glycoprotein-complex (DGC) in muscle cells, dystroglycan provides a critical anchor between the ECM and the actin cytoskeleton to protect the sarcolemma from the forces of muscle contraction (3, 4). Similarly, dystroglycan expressed on astrocyte endfeet in the brain links to the basement membrane of endothelial cells lining blood vessels to help maintain the blood brain barrier (5). Thus, loss of functional dystroglycan or breaking of the mechanical link it provides is generally detrimental to cells. Whole animal dystroglycan deletion leads to embryonic lethality (6), and conditional knockouts in muscle and brain tissues result in drastic tissue disorganization (7, 8). Moreover, mutation or deletion of glycosyltransferases that post-translationally modify dystroglycan breaks the link between dystroglycan and the ECM, and is associated with subtypes of muscular dystrophy, cancer, and brain defects (9–11). Similarly, mutations and truncations of dystrophin, the adaptor protein that connects the intracellular portion of dystroglycan to actin, lead to Duchenne Muscular Dystrophy (12–14).

However, dystroglycan’s link to the ECM must be dynamically modulated in contexts where cellular plasticity is required, such as cell migration, wound repair, or function of neural synapses. One such mechanism of modulation of dystroglycan’s adhesion to the ECM is post-translational cleavage by matrix metalloproteases (MMPs) just outside of the transmembrane domain. However, the mechanism of regulation of MMP cleavage of dystroglycan has not been investigated, despite several lines of evidence to suggest a role for abnormal proteolysis in disease pathogenesis. Proteolysis of β-dystroglycan to a 31-kDa fragment is enhanced in several pathogenic states, such as cancer (15–17), muscle diseases (18, 19), and autoimmune disorders (5). Additionally, high levels of MMPs are found in biopsies of muscular dystrophy patients (20) and muscular dystrophy phenotypes are ameliorated in mice treated with metalloprotease inhibitors (21–23). Moreover, MMP-2/MMP-9 double-knockout mice in which MMP cleavage of dystroglycan is blocked are resistant to the onset of autoimmune encephalomyelitis, underscoring the relevance of this cleavage in disease pathogenesis (5).

Dystroglycan (DG) is encoded by the *Dag1* gene and translated as a single polypeptide precursor that undergoes post-translational processing to generate two non-covalently interacting subunits at the cell-surface: the extracellular and highly glycosylated α-DG that binds laminin and other ligands in the ECM and the transmembrane β-DG that binds to the cytoskeleton via adaptor proteins such as dystrophin (24, 25). The non-covalent association of α-and β-DG occurs within a putative Sea urchin-Enterokinase-Agrin-like (SEA-like) domain, located extracellularly between the ECM-binding and transmembrane regions (26). Importantly, this SEA-like domain also houses the predicted MMP cleavage site(s) and harbors one of the two patient-derived muscular dystrophy mutations (C669F) identified in dystroglycan thus far (27). Intriguingly, the SEA-like domain bears striking structural homology to the heterodimerization domain (HD) of Notch receptors, which is also non-covalently associated as a result of furin cleavage during maturation and houses Notch’s critical ADAM protease site that is exposed by intercellular forces to trigger Notch activation. In Notch, the neighboring N-terminal Lin-12-Notch (LNR) domain and the HD domain cooperate to form the Negative Regulatory Region (NRR) which conformationally protects the cleavage site from its protease until ligand engagement (28–31).

Motivated by the incidence of abnormal proteolysis of dystroglycan in pathogenic contexts, the putative structural homology to the NRR of Notch, and the presence of disease-related mutations in this domain (27, 32–36), we present here a study of the regulation of proteolysis in the extracellular juxtamembrane region of dystroglycan. Through a series of biochemical and cellular assays, we show that dystroglycan proteolysis is conformationally regulated. Furthermore, muscular dystrophy mutations disrupt the integrity of the proteolysis domain sensitizing dystroglycan to proteolysis. Importantly, increased proteolysis correlates with increased rates of cell migration in wound healing assays, linking proteolysis to a disease-relevant cellular phenotype. Finally, we find that several cancer mutations that map to dystroglycan’s proteolysis domain similarly lead to increases in proteolysis and rates of cell migration, potentially revealing a new pathogenic mechanism in cancer.

## Results

### Dystroglycan α/β-processing requires both Ig-like and SEA-like domains

In order to understand how dystroglycan proteolytic cleavage is regulated, we first aimed to map the boundaries of the extracellular juxtamembrane domain that houses both the putative MMP cleavage site(s) ~25 amino acids from the membrane and the α/β-processing site defining the α/β dystroglycan transition located ~50 amino acids from the membrane (19, 37). Previous studies of mucin SEA domains have revealed that the post-translational autoproteolytic α/β-processing event can only occur if the domain is correctly folded, due to the necessity of a precisely strained loop to permit the serine in the cleavage sequence to nucleophilically attack the neighboring glycine (38). Thus, we utilized SEA-domain α/β-processing as an indication of correct protein folding to map the minimal functional unit of dystroglycan’s proteolysis domain and devised a series of constructs including the SEA-like domain and its neighboring Ig-like domain. Dual-epitope tagged full-length and truncation constructs of dystroglycan’s proteolysis domain were designed using predicted secondary structure as a guide and secreted into conditioned media of HEK293T cells (Fig 1A). The construct also included a C-terminal Fc-Fusion tag to enable immunoprecipitation with Protein A beads.

**FIG 1.**
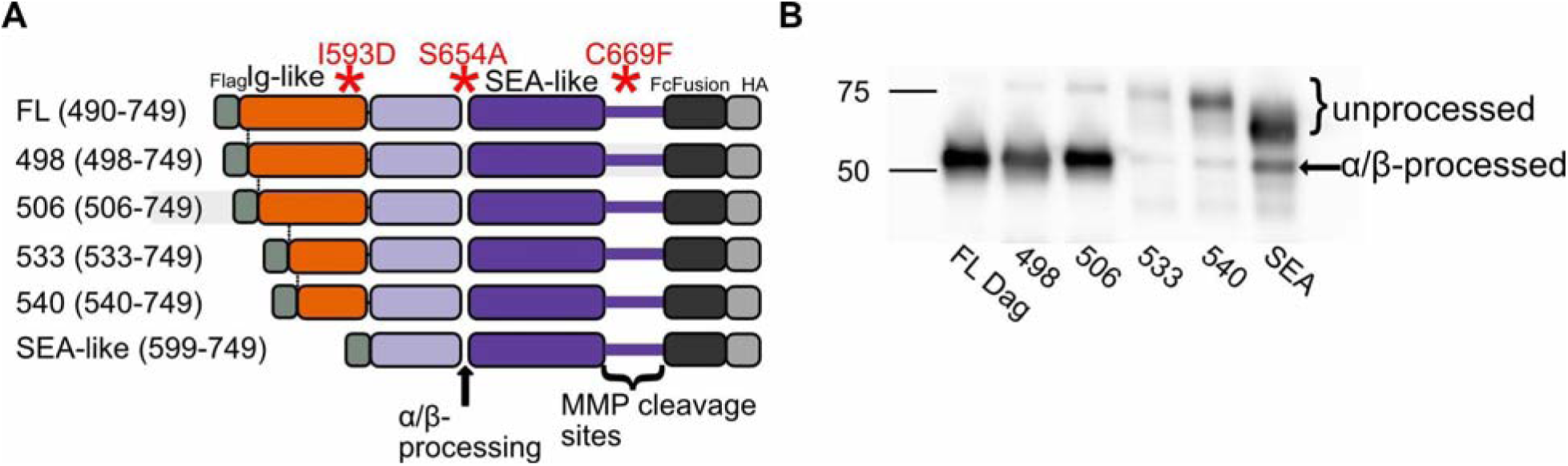
Dystroglycan α/β-processing requires both Ig-like and SEA-like domains (A) Infographic of constructs used to map dystroglycan proteolysis domain. Constructs comprised of Ig-like (orange) and SEA-like (purple) domains contain N-terminal FLAG and C-terminal HA epitope tags as well as a C-terminal Fc-fusion. Sites of α/β-processing, MMP cleavage, and disease-related mutations (red asterisks) are depicted (B) α-HA western blot comparing unprocessed and α/β-processed protein levels in dystroglycan truncation constructs from A. secreted into conditioned media and immunoprecipitated with Protein A beads.

The protein was enriched from the conditioned media with protein A beads and Western blotted for the C-terminal HA tag. Two potential α-HA-tagged protein species were expected: a band of ~50 kDa corresponding to the correctly α/β-processed C-terminus for all of the constructs and a variable-sized larger band corresponding to the full-length unprocessed protein. The construct containing the entire Ig-like domain in addition to the SEA-like domain showed a dominant 50 kDa band and faint 75 kDa band, reflecting >90% α/β-processing. (Fig 1B). However, the N-terminal truncation constructs exhibited an increasing ratio of unprocessed to processed dystroglycan, and the construct containing solely the SEA-like domain showed the lowest degree of α/β-processing. These data suggest that the Ig-like domain folds in concert with the SEA-like domain to support dystroglycan α/β-processing. We will term the portion of dystroglycan comprised of the full Ig-like and SEA-like domains the dystroglycan “proteolysis domain.”

### MMP cleavage of the dystroglycan proteolysis domain is conformationally regulated

We next wanted to ask whether the MMP cleavage, predicted to occur ~20 amino acids external to the transmembrane domain (37, 39), was conformationally regulated in the context of the isolated proteolysis domain, as in Notch receptors. To study the accessibility of dystroglycan’s proteolytic site(s) to exogenous MMPs, dual-epitope tagged full-length and SEA-like domain constructs used in the mapping studies were expressed and secreted from HEK-293T cells. The immunoprecipitated dystroglycan constructs were incubated with increasing amounts of activated MMP-2 and MMP-9 for 1 hour, eluted from the beads, and Western blotted against the HA tag. Upon addition of exogenous MMPs, the full-length dystroglycan proteolysis domain remains a single band on the α-HA western blot while a faster-migrating species appears for the SEA-like domain alone construct (Fig 2A). This suggests that the MMP site is exposed when the Ig-like domain is not present, likely due to incorrect folding of the domain, which is consistent with the truncation studies in Fig 1B.

**FIG 2.**
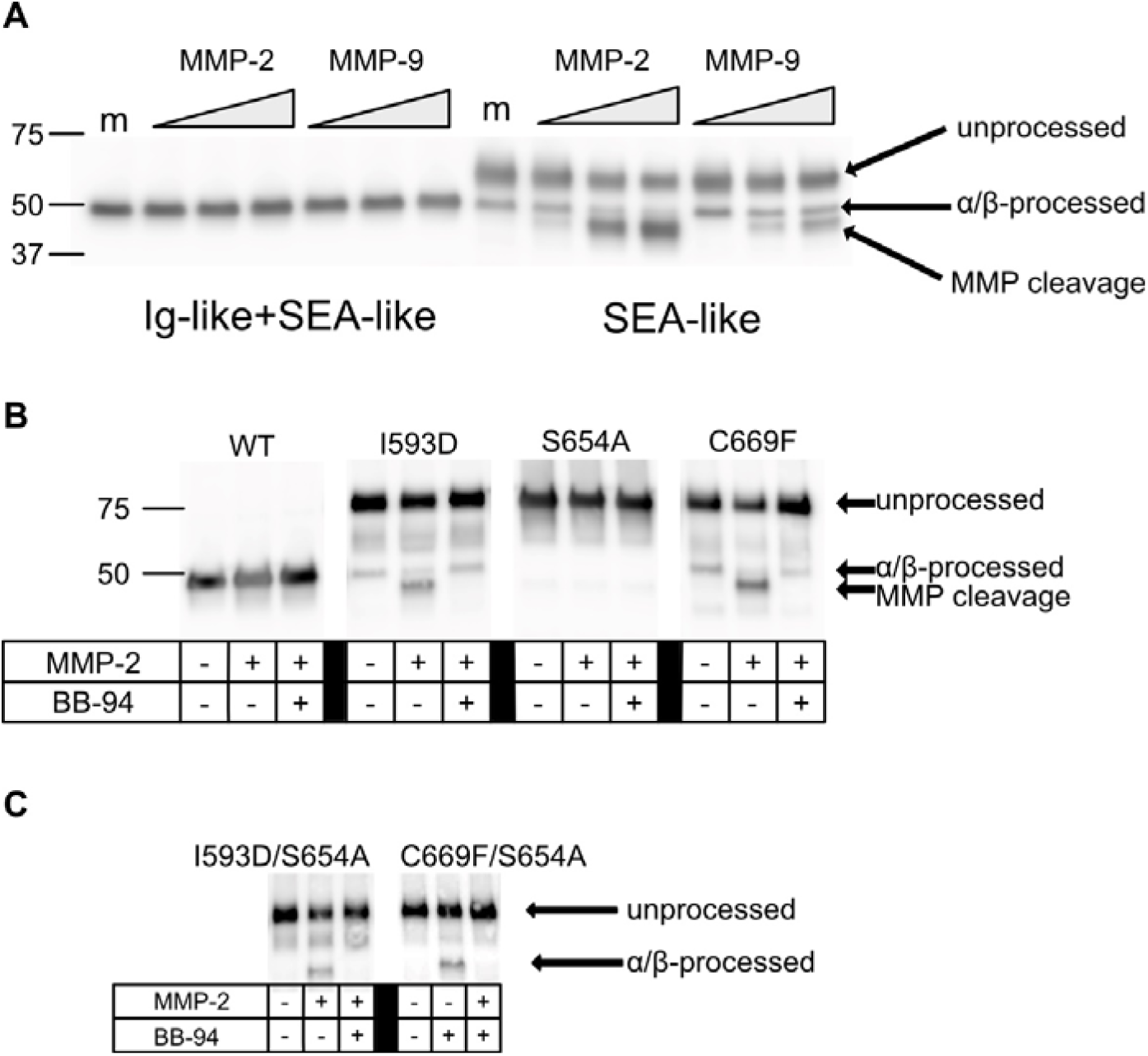
MMP cleavage of the dystroglycan proteolysis domain is conformationally regulated. (A) α-HA western blot of full length dystroglycan proteolysis domain (Ig-like and SEA-like domains) versus SEA-like domain only. Increasing concentrations of activated MMP-2 and MMP-9 were reacted in MMP buffer with protein secreted from conditioned media and immunoprecipitated with Protein A beads. m= MMP buffer only. (B) α-HA western blot of secreted dystroglycan proteolysis domains harboring disease mutations. Reactions were performed in MMP buffer and 0.73 μg/ml activated MMP-2 and 40 μM BB-94 were added as noted. (C) α-HA western blot of secreted dystroglycan proteolysis domains harboring disease mutations in the context of blocked α/β-processing (S654A mutation). Constructs were secreted from HEK293T cells and enriched using Protein A beads. Reactions were performed in MMP buffer and 0.73 μg/ml activated MMP-2 and 40 μM BB-94 were added as noted.

Since protection of the MMP site in the context of isolated proteolysis domains relies on the correct conformation of the domain, we next wanted to test whether mutations associated with muscle disease found in humans and model organisms alter the conformation of the domain and expose the protease site. To date, three disease-related mutations within the dystroglycan proteolysis domain are associated with muscular dystrophy, but how they contribute to disease pathogenesis is unclear. I593D, a mutation located in the Ig-like domain, leads to a muscular dystrophy phenotype in zebrafish (40). The S654A mutation blocks α/β-processing and has been observed to cause a muscular dystrophy phenotype in mice (33). Finally, the C669F mutation located near the C-terminus of the SEA-like domain, which disrupts a putative disulfide linkage, was discovered in patients with a severe form of muscular dystrophy (27). It should first be noted that decreased α/β-processing is observed for the three mutations (Fig 2B). While expected for the S654A mutant in which the cleavage site is abolished, the incorrect processing of C669F and I593D suggests that they disrupt the structure of the domain, much like the truncation constructs described in Figures 1B and 2A. Indeed, addition of exogenous MMPs to proteolysis domains harboring the I593D and C669F mutations resulted in the appearance of an MMP-cleaved species that was abrogated by the addition of batimastat (BB-94), a global metalloprotease inhibitor (Fig 2B). In contrast, previous structural work of mucin SEA domains has shown that mutating the α/β-processing cleavage site as in S654A does not impact the overall fold of the SEA domain (38). Thus, the S654A cleavage mutant is not cleaved by MMPs, suggesting an intact proteolysis domain. Similar levels of MMP cleavage were observed in double mutants where α/β-processing was fully blocked by creating I593D/S654A and C669F/S654A constructs (Fig 2C).

### Proteolysis to the 31-kDa β-dystroglycan cleavage fragment also depends on the structural integrity of the proteolysis domain

We next asked whether the MMP resistance phenotype of the wild-type proteolysis domain and the proteolysis sensitive phenotype observed in truncation and muscular dystrophy disease mutants was recapitulated in the full-length receptors. Full-length dystroglycan constructs (Fig 3A) were transiently transfected into Cos7 cells (Fig 3B), HEK293T, and U2OS cells (Fig 3C). Cell lysates were Western blotted for β-dystroglycan. Full-length wild-type dystroglycan undergoes normal α/β-processing and no appreciable lower molecular weight species are observed, as expected. In contrast, the I593D and C669F mutations both disrupt α/β-processing and a new 31 kDa fragment appears, which corresponds to the size of β-dystroglycan that is cleaved by MMPs. To confirm that this band corresponds to a proteolytic fragment of the correct size, we expressed a construct termed “39-end” beginning 39 amino acids outside of the transmembrane domain, approximately corresponding to the expected MMP-cleaved species. This construct migrated similarly to the observed 31 kDa fragment. To further test that the new species corresponds to proteolytic events near the C-terminus of the SEA-like domain and is not just an artifact of overexpression, we created a construct Δ39 in which the putative MMP cleavage sites were deleted (removal of amino acids 711–749 directly before the transmembrane domain). When this deletion is present in the context of the C669F mutation, the 31 kDa species is abolished (Fig 3B). Moreover, in the context of a construct termed “ΔIgΔSEA” that removes all of the proteolysis domain except amino acid 711–749 in which one might expect constitutively exposed proteolysis sites, a large fraction of β-dystroglycan is further proteolyzed into the 31 kDa species.

**FIG 3.**
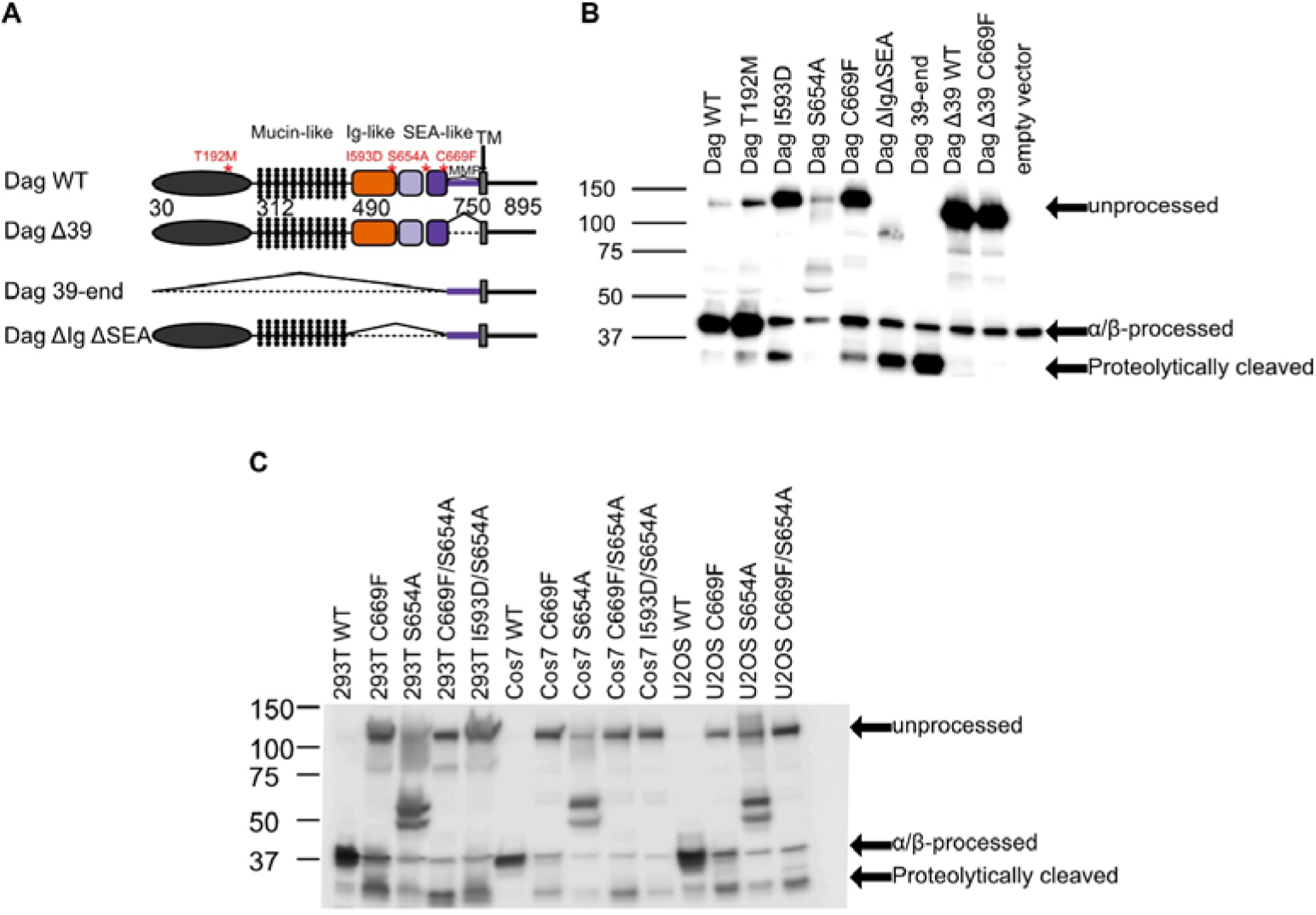
Presence of the 31-kDa B-dystroglycan proteolytic fragment depends on the structural integrity of the proteolysis domain. (A) Schematic of full-length dystroglycan constructs. Red asterisks denote location of disease mutations. Dashed lines denote domain deletions. (B) β-dystroglycan western blot of full-length dystroglycan constructs transiently transfected in Cos7 cells. Full-length β-dystroglycan migrates at 42 kDa while proteolytically-cleaved β-dystroglycan migrates at 31 kDa. T192M, I593D, and C669F disease-related mutations and deletion of the entire proteolysis domain save putative protease sites (ΔIgΔSEA) show a 31 kDa proteolysis product. Deletion of protease sites Δ39 in the context of C669F abolishes 31 kDa band. The 39-end construct provides an approximate size marker for the expected proteolytic fragment. (C) β-dystroglycan western blot of full-length dystroglycan constructs transiently transfected in HEK293T, Cos7, and U2OS cells. Full-length β-dystroglycan migrates at 42 kDa while proteolytically-cleaved β-dystroglycan migrates at 31 kDa. I593D, S654A, and C669F are disease-related mutations.

### Modulation of Dystroglycan proteolysis alters cell migration

Thus far, we have observed that proteolysis of dystroglycan depends on the structural integrity of the “proteolysis domain.” Wild-type receptors and receptors where putative cleavage sites have been deleted are resistant to cleavage while domain truncations and disease-related mutations sensitize dystroglycan to proteolytic cleavage. To determine if this proteolysis is relevant to dystroglycan’s critical roles in the cell, we next asked whether modulating dystroglycan proteolysis results in cellular phenotype changes related to dystroglycan’s role as an adhesion receptor. Dystroglycan mutations/truncations tested for proteolysis in the western blot were used to modulate levels of proteolysis and determine effects on cell migration in a wound-healing assay. We used two cell lines known to produce ECMs containing the major dystroglycan ligand laminin during normal cell culture conditions. Constructs were transiently transfected into 3T3-L1 fibroblast cells and U251 glioblastoma cells (Fig 4), and a scratch was introduced after 24 hours. Migrating cells were imaged at different time points up to 18 hours post-scratch, and the wound area was calculated. Analysis of three areas of the wound in duplicate wells at each time point were performed and averaged, and consistent results were obtained over at least four separate experiments performed on different days. Representative images of wounds for Dag WT, Dag + BB-94 and Dag C669F in U251 cells are shown in in Figure 4A. Over-expression of wild-type dystroglycan in 3T3 cells modestly slowed down cell migration rates (Fig 4B). Treating the wild-type-overexpressing cells with BB-94 significantly slowed down wound-healing, suggesting that broadly blocking proteolytic cleavage in cells alters the balance of adhesion versus migration to favor adhesion. On the other hand, cells transfected with disease-related mutations C669F, I593D, and the truncation mutant ΔIgΔSEA migrated at a significantly faster rate than wild-type, suggesting that the alterations of the proteolysis domain that enhance proteolysis favor cellular migration at the expense of adhesion. Treating C669F expressing cells with BB-94 rescued the enhanced migration phenotype, as expected.

**FIG 4.**
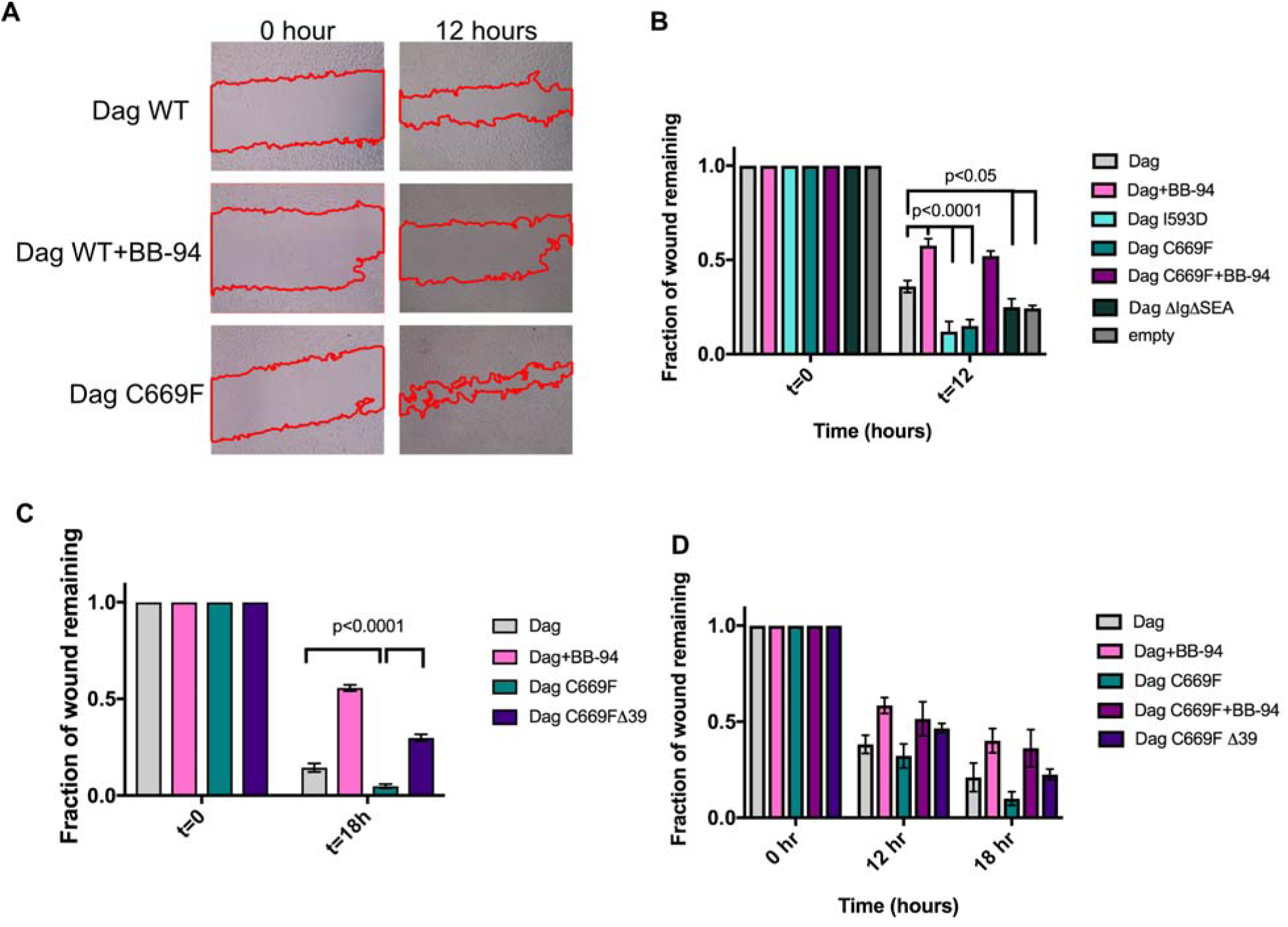
Modulation of dystroglycan proteolysis alters cell migration. (A) Representative brightfield images of wounds scratched in U251 cells transiently transfected with dystroglycan constructs at t=0 hours and t=12 hours. Red outline denotes wound edges. (B-D) Bar graphs denoting fraction of wounds remaining for dystroglycan constructs or empty vector at different time points. Wound areas were calculated for 8–12 images for each construct at each time point, averaged, and normalized to wound area at t=0. Error bars represent the SEM of 8–12 measurements, and statistical significance was determined with a two-way ANOVA followed by a post hoc Bonferroni test in Graphpad Prism. (B, C) Wound healing assays performed on different days in 3T3 cells. (D) Wound healing assay perfromed in U251 cells.

Since global metalloprotease inhibitors are not specific and are likely blocking proteolysis of other receptors in the cells, we also tested Dag Δ39 constructs in which the putative MMP sites are deleted. The deletion of the putative proteolytic cleavage sites rescues the effect of the C669F mutation on migration, as cells migrate similarly to wild-type (Fig 4C). This result suggests that proteolytic cleavage of C669F near the C-terminus of the proteolysis domain alters a cell’s migration phenotype to favor faster migration/less adhesion, linking enhanced proteolysis to a disease-relevant cellular phenotype. The migration phenotypes were consistent in U251 glioblastoma cells (Fig 4E).

### Cancer mutations in dystroglycan’s proteolysis domain enhance proteolysis and alter cell migration

Thus far we have shown that muscle disease-related mutations in the proteolysis domain enhance dystroglycan cleavage to the 31-kDa fragment which correlates with faster cell migration. A faster cell migration phenotype has also been observed in the context of hypoglycosylation/reduced laminin binding of dystroglycan in the context of cancer (9). This suggests that proteolysis may provide an alternative mechanism relevant to disease pathogenesis for breaking the dystroglycan mechanical link. We were thus inspired to test whether cancer mutations within dystroglycan’s proteolysis domain exhibit abnormal proteolysis/migration akin to dystroglycan glycosylation defects. The COSMIC database reports 123 missense mutations of dystroglycan, and the FATHMM algorithm (41) predicts that 120 mutations are pathogenic. 19 of these map to the proteolysis domain. We selected mutations from the COSMIC database that spanned the Ig-like and SEA-like domains and were flagged as “pathogenic” by FATHMM prediction (Table 1).

**Table 1.** Disease-related mutations used in this study

| Mutations studied | Location in proteolysis domain | Disease context | $\alpha/\beta$-processing | Extent of proteolysis to 31 kDa | Rate of wound healing | Reference |
| --- | --- | --- | --- | --- | --- | --- |
| WT | - | - | +++ | + | + |  |
| Y575H | Ig-like | Adenocarcinoma | +++ | +++ | +++ | generated by the TCGA Research Network |
| I593D | Ig-like | Muscular Dystrophy | + | ++++ | +++ | (40) |
| R640Q | SEA-α | Adenocarcinoma | +++ | +++ | +++ | (35) |
| S654A | Autoproteolysis site | Muscular Dystrophy | none | + |  | (33) |
| C669F | SEA-β | Muscular dystrophy | + | ++++ | +++ | (27) |
| G676V | SEA-β | Glioblastoma | +++ | ++ | ++ | generated by the TCGA Research Network |
| S678G | SEA-β | Malignant melanoma | +++ | + | + | (36) |
| D738H | SEA-β | Diffuse large B cell lymphoma and Adenocarcinoma | +++ | + | +++ | (34) |
| R739W | SEA-β | Glioblastoma | +++ | | ++ | generated by the TCGA Research Network |

We first transfected full-length dystroglycan constructs harboring missense mutations into Cos7 cells to detect the 31-kDa proteolytic fragment. Interestingly, unlike the muscular dystrophy mutations, all of the cancer mutants displayed normal α/β-processing, suggesting that these mutations do not disrupt the fold of the proteolysis domain to the same extent as the muscular dystrophy related mutations (Fig 5A). Comparing the ratio of the 31 kDa proteolysis fragment to the sum of unprocessed and processed β-dystroglycan bands to calculate fraction proteolyzed reveals that most of the cancer mutants enhance proteolysis compared to wild-type dystroglycan, though not to the same extent as the muscle disease C669F/I593D mutations. We next tested a subset of these mutations, particularly the mutations with enhanced proteolysis, in the wound healing assay. Indeed, most of the mutations tested resulted in statistically significant increases in the rate of wound closure (Fig 5B), suggesting that enhanced proteolysis of dystroglycan via mutation of the proteolysis domain may represent a new pathogenic mechanism in cancer, with similar effects to breaking the dystroglycan link via hypoglycosylation (9, 42, 43).

**FIG 5.**
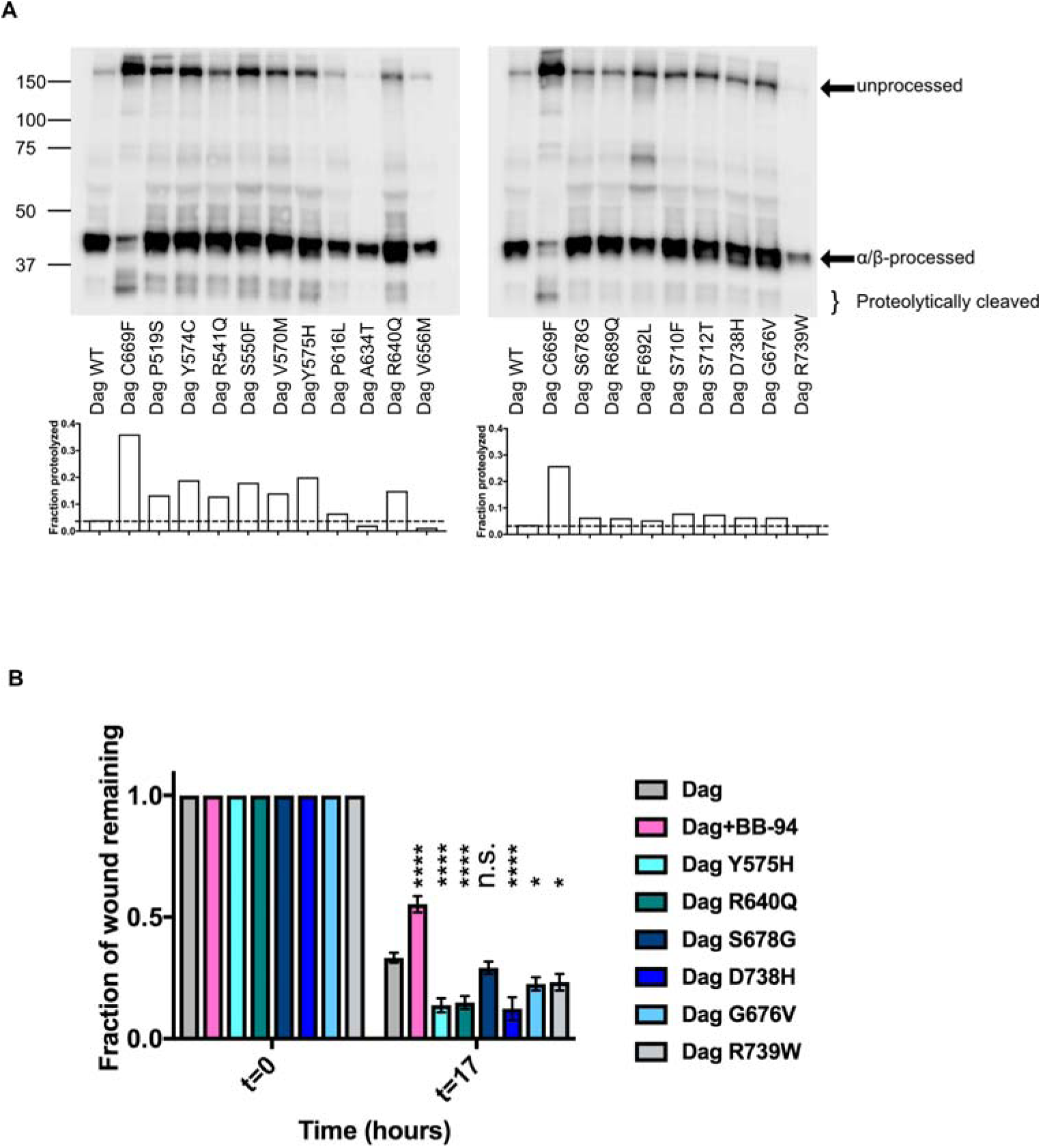
Cancer mutations in dystroglycan proteolysis domain enhance proteolysis and wound healing. (A) β-dystroglycan western blot of full-length dystroglycan constructs transiently transfected in Cos7 cells. Full-length β-dystroglycan migrates at 42 kDa while proteolytically-cleaved β-dystroglycan migrates as a doublet at 31 kDa. Fraction proteolyzed was calculated by taking the ratio of the 31-kDa band and the sum of the processed and unprocessed bands using gel quantitation features in ImageJ. (B) Several cancer mutations were evaluated in a wound healing assay of constructs transiently transfected in 3T3 cells. Bar graphs denoting fraction of wounds remaining for dystroglycan constructs at different time points. Wound areas were calculated for 9 images for each construct at each time point, averaged, and normalized to wound area at t=0. Error bars represent the SEM of 9 measurements, and statistical significance was determined with a two-way ANOVA followed by a post hoc Bonferroni test in Graphpad Prism. Statistical significance of fraction of wound healed at a given condition with respect to wild-type is depicted as **** (p<0.0001), * (p<0.05) or n.s. p> 0.05.

## Discussion

The adhesion receptor dystroglycan provides a critical mechanical link between the ECM and the actin cytoskeleton, most notably in muscle and brain cell types. In muscle, dystroglycan is at the core of the multi-protein DGC, acting as a shock-absorber to protect muscle cells from contractile forces. In the brain, dystroglycan organizes laminin in the basal lamina to maintain the blood brain barrier as well as provide the cortical infrastructure necessary for neuronal migration. Disruption of the mechanical link via impairment of dystroglycan binding to the ECM or to the actin-adaptor dystrophin is associated with diseases such as muscular dystrophy and cancer (1). The ECM to cytoskeleton anchor that dystroglycan provides can also be disrupted by proteolytic cleavage of dystroglycan by MMPs just outside of the transmembrane domain, and several lines of evidence suggest a role for abnormal dystroglycan proteolysis in disease pathogenesis. Thus our study aimed to shed light on how proteolysis is regulated in order to provide critical insights into dystroglycan’s multiple functions as a mechanical link and adhesion receptor in the cell and to provide insights into disease pathogenesis.

The first conclusion of this work is that dystroglycan proteolysis by MMPs is conformationally regulated by a juxtamembrane “proteolysis domain”, analogous to regulation of proteolysis in Notch receptors. First, we find that dystroglycan’s SEA-like domain, which harbors both the α/β-processing site and the putative MMP cleavage sites, cooperates with its adjacent Ig-like domain to support the non-covalent complex formed from α/β-processing during receptor maturation. Autoproteolysis, which depends on correct folding of the domain, does not occur in the SEA domain alone and only occurs in the presence of its neighboring Ig-like domain. While MUC1 (45) and MUC16 (46) SEA domains and the SEA-like domain of the receptor tyrosine phosphatase IA-2 (47) exist in the absence of neighboring domains, the dystroglycan SEA-like domain appears to be more similar to the SEA-like domains of Notch (29, 30, 48) and EpCAM (49), which extensively interact with neighboring N-terminal domains. We find that the dystroglycan proteolysis domain in isolation is resistant to exogenous metalloproteases, but becomes susceptible to proteolysis when the domain is truncated or encodes muscle disease-related mutations, much like in Notch receptors (Fig 6) (44, 50). A similar trend is observed in full-length receptors; low levels of the 31 kDa fragment are observed for the wild-type receptor, but levels are dramatically increased for constructs in which a majority of the proteolysis domain has been removed and for receptors encoding disease-related mutations. Moreover, the aberrant proteolysis in the C669F mutant is abrogated when the putative metalloprotease sites are deleted from the construct.

**FIG 6.**
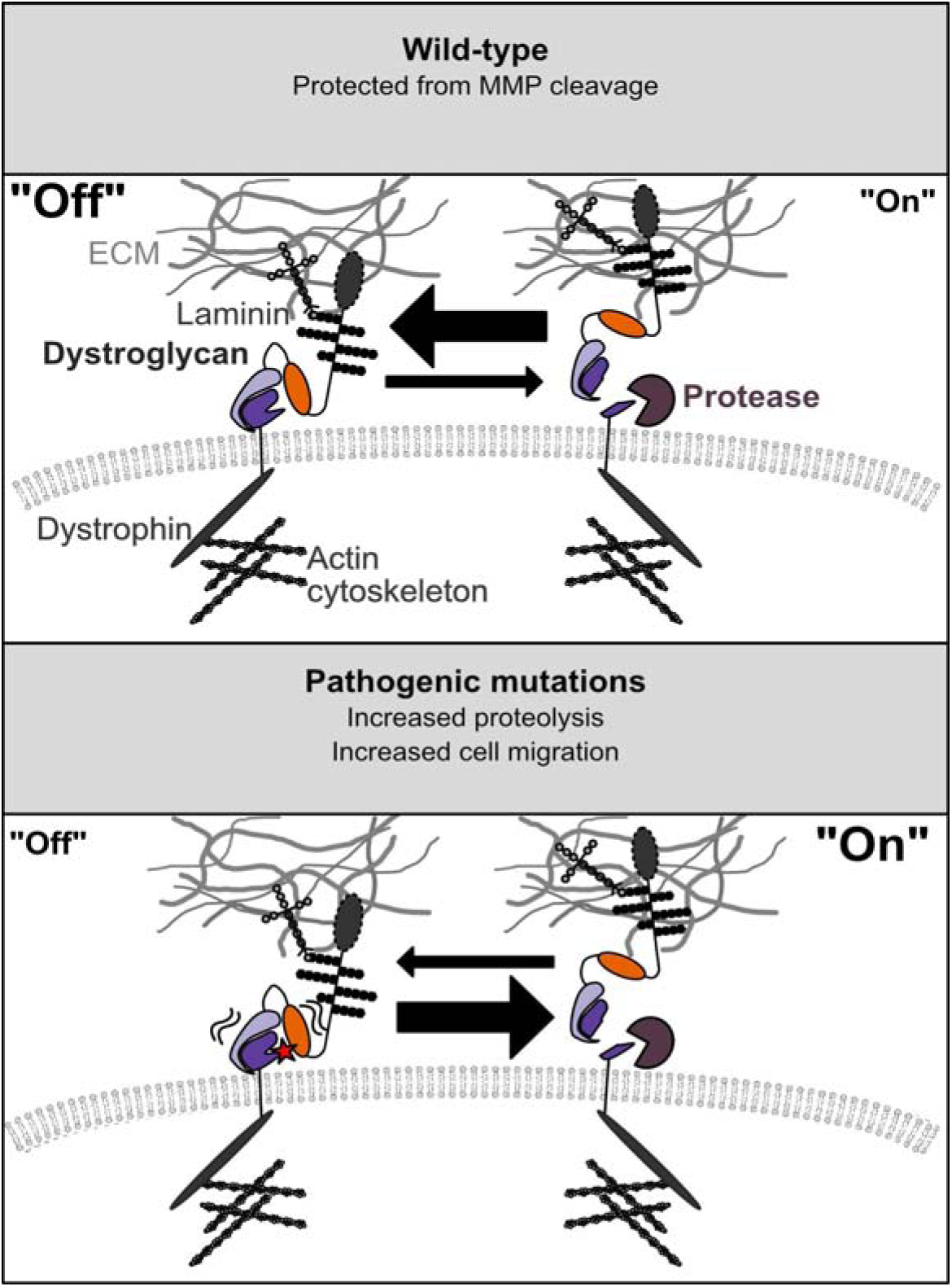
Model for conformational control of dystroglycan proteolysis in wild-type and mutated dystroglycan. Wild-type dystroglycan provides an important mechanical connection between the ECM and actin cytoskeleton. Mutations, such as C669F, destabilize the proteolysis domain, rendering it sensitive to proteolysis and increasing cell migration rates. Cellular factors characterizing pathogenic states, such as increased ECM stiffness and increased MMPs, may also increase dystroglycan proteolysis rates and contribute to pathogenicity.

The second conclusion of this work is that mutations in the dystroglycan proteolysis domain destabilize the domain, resulting in sensitization of the proteolysis domain to proteolytic cleavage and thus disruption of the critical ECM to actin tether (Figure 6). Only two mutations in the DAG1 gene have been reported in patients with muscular dystrophy; the T192M mutation near the dystroglycan N-terminus, which leads to hypoglycosylation and thus deficits in ECM binding (32) and the C669F proteolysis domain mutation reported in two patients (27), with unknown mechanism of disease pathogenesis. We also examined previously uncharacterized dystroglycan mutations mined from cancer data sets in COSMIC and TCGA (Table 1 The muscle disease related mutations disrupt α/β-processing, suggesting that they drastically alter the fold of the SEA-like domain and expose the normally occluded protease sites. Isolated proteolysis domains harboring these mutations are cleaved by exogenous MMPs, and a large fraction of the receptor is cleaved into the 31 kDa species in the context of the full-length receptor. On the other hand, the cancer mutations we investigated generally exhibit normal α/β-processing, suggesting that the structure of the SEA-like domain is not drastically altered. Further work is need to explore the mechanism of proteolytic exposure in the cancer mutants, but the observed graded structural disruption is similar to observations of leukemia-derived mutations found in the NRR of Notch receptors (44). Nevertheless, mutations of both classes result in enhanced proteolysis in the context of the full-length receptor that correlates with increased cellular migration rates.

Finally, the third conclusion of this work is that proteolysis of dystroglycan by MMPs outside of the transmembrane region correlates with faster migration in cells. Exploiting the tunable proteolytic sensitivity of the constructs used in this work, we found that mutants that lead to increased proteolysis result in faster migration phenotypes while mutants that block proteolysis are linked to slower migration phenotypes similar to effects of pan-metalloprotease inhibitors. This effect was observed in both human fibroblast and glioblastoma cells, which are known to produce ECM laminin when cultured (51, 52). The finding of an increased migration phenotype when proteolysis disrupts the dystroglycan mechanical link is similar to previous studies in cancer cell lines in which increased cell migration was observed when dystroglycan was hypo-glycosylated and thus defective in binding the ECM in several cancer cell lines (9) or when an antibody was used to block dystroglycan binding to laminin in prostate cancer cell lines (42). Interestingly, hypo-glycosylated and/or reduced dystroglycan expression has been associated with more metastatic cancers (9, 16, 53, 54). Moreover, patients harboring the C669F mutation, which has been observed in two siblings (27), exhibit severe muscle-eye-brain phenotypes which is characterized by neuronal migration defects, providing another link between the observed cellular phenotype and disease pathogenesis. On the other hand, overexpression of dystroglycan in breast cancer cells reduces tumorigenicity (55), hinting that restoring broken dystroglycan links could be a therapeutic option. Indeed re-expressing deficient glycosyltransferases, which restores dystroglycan linkage to the ECM and rescues the fast migration phenotype in glycosylation-deficient dystroglycan contexts, is being pursued as a current therapeutic avenue (56)

Together, our findings that proteolysis is normally controlled by the conformation of the domain, that mutations destabilize the domain and aberrantly increase proteolysis, and that enhanced proteolysis is correlated with a cellular phenotype strongly linked to disease pathogenesis, suggest that blocking dystroglycan proteolysis may be a valuable therapeutic avenue. Indeed, treatment of muscular dystrophy mouse models with protease inhibitors ameliorates symptoms. However, our study provides a rationale and platform for developing specific inhibitors of dystroglycan proteolysis. Since dystroglycan proteolysis is conformationally controlled as in Notch receptors, it may be possible to develop conformation-specific antibodies that clamp the domain in a closed conformation to prevent proteolysis, a strategy which has been successfully employed in Notch receptors (57, 58). The results of this work also beg further investigations into the cellular stimuli leading to aberrant proteolysis of dystroglycan in cases where the receptor is not mutated.

## Acknowledgements

We would like to thank the Odde lab at the University of Minnesota for the U251 cells and the Titus lab for the COS7 cells. We would like to thank Jim Ervasti and Kevin Campbell for helpful discussions and Eric Aird for useful comments on this manuscript. This study was supported by an NIH NIGMS R35GM119483 grant to WRG. ANH received salary support from AHA Fellowship 17PRE33330000 and is an ARCS Fellow. WRG is a Pew Biomedical Scholar.

## Author contributions

ANH and WRG designed experiments. AH acquired data. AH and WRG analyzed and interpreted data. AH and WRG wrote and revised the manuscript.

## Experimental procedures

### Materials

Antibodies that were used in this study-Flag antibody: Sigma, HA antibody: Covance, β-Dystroglycan antibody against 15 amino acids at the extreme C-terminus: Leica Biosystems. Restriction enzymes were purchased from New England Biolabs (NEB). Cloning was performed using In-Fusion HD Cloning Mix (Clontech). All electrophoresis supplies were purchased from Bio-Rad. Dual Luciferase kit was purchased from Promega. MMP-2 and MMP-9 were purchased from R&D Systems. Mutations within dystroglycan’s proteolysis domain were created using Pfu Turbo Polymerase (Agilent) and DpnI (NEB).

### Construct creation

Full length human dystroglycan cDNA in CMV-6 plasmid was obtained from Origene and used as a template for all of the cloning.

For expression of secreted proteolysis domain: Ig-like+SEA-like domain (amino acids 490–749) and SEA-like domain (599–749) constructs were cloned into pcDNA3 vector containing an N-terminal Notch signal sequence and Flag tag and C-terminal Fc-fusion and HA tags.

### Mammalian cell culture

All of the cells used in this paper were cultured in Dulbecco’s Modified Eagle Medium (DMEM, Corning) supplemented with 10% FBS, 100 units/mL penicillin, and 100 units/mL streptomycin. Cos7 cells were a kind gift from Dr. Margaret Titus. U251 cells were a kind gift from Dr. David Odde. U2OS and HEK293T cells were purchased from ATCC. MS5 and MS5-DLL4 cells were a kind gift from Steve Blacklow.

All transfections were performed in Opti-MEM (Gibco) with Lipofectamine 3000 (Invitrogen) according to manufacturer instructions.

### MMP activation

MMP-2 and MMP-9 were activated using p-aminophenylmercuric acetate (APMA) in MMP assay buffer (50 mM Tris pH 7.5, 10 mM CaCl_2_, 150 mM NaCl, 0.05% Brij-35 (w/v)) according to manufacturer instructions. MMP-2 was activated for 1 hour at 37°C. MMP-9 was activated for 24 hours at 37°C according to manufacturer instructions.

### MMP assay

10 cm plates of HEK-293T cells were transiently transfected with dystroglycan proteolysis domain. The conditioned media was collected 48 hours post-transfection. Conditioned media was bound to magnetic protein A beads and incubated for 1 hour at 4°C. The beads were washed thoroughly with wash buffer (50 mM Tris pH 7.5, 150 mM NaCl, 0.1% NP-40). The beads were split into separate tubes, MMP buffer was added (50 mM Tris, 10 mM CaCl_2_, 150 mM NaCl, 0.05% Brij-35 (w/v), pH 7.5) and activated MMP and BB-94 added as appropriate. For the experiment using increasing amounts of MMP in Figure 2A: 0.07 μg/mL, 0.37 μg/mL, and 0.73 μg/mL of respective MMPs were added, and the tubes were incubated for 1 hour at 37°C. The remaining protein was eluted by incubating the beads with 2x SDS sample buffer with 100 mM DTT for 5 minutes at room temperature.

### Western blot

Lysates and dystroglycan proteolysis domain were run on a 4–20% SDS-PAGE gel including 2 mM sodium thioglycolate in the running buffer. The protein was then transferred to a nitrocellulose membrane using a Genie Blotter (Idea Scientific) and blocked with 5% milk. The antibodies were diluted 1:1000 in 5% bovine serum albumin (BSA). A goat-anti mouse HRP conjugated antibody (Invitrogen). Western blots were imaged using chemiluminescent buffer (Perkin Elmer Western Lightning Plus ECL) and the Amersham 600UV (GE) with staff support at the University of Minnesota-University Imaging Center.

### Immunofluorescence (if we need for supplement)

U2OS cells were transiently transfected with Notch chimera constructs for 24 hours. Then, the cells were incubated with 1:500 Flag APC (Invitrogen) antibody in D10 for 45 minutes at 37°C. The cells were washed with PBS and imaged in clear DMEM on an EVOS-FL Auto microscope.

### Wound healing assay

3T3 cells were reverse transfected in duplicate in a 24 well dish with dystroglycan constructs or an empty vector. After 36 hours, a scratch was made with a P200 pipette, the cells were washed once with PBS, and placed in DMEM supplemented with 5% FBS. Images were taken at respective time points using an EVOS FL Auto microscope. Four image areas were marked and monitored for each well, and wound area measurements were performed independently with Adobe Photoshop and the MRI Plugin for ImageJ. The data from four independent experiments were analyzed and plotted using Prism. Each wound area was normalized where 100% of the area was at time=0. The normalized data was then used for a two-way ANOVA analysis to determine significance.

